# Dramatic HIV DNA degradation associated with spontaneous HIV suppression and disease-free outcome in a young seropositive woman following her infection

**DOI:** 10.1101/651752

**Authors:** Philippe Colson, Catherine Dhiver, Catherine Tamalet, Jeremy Delerce, Olga O. Glazunova, Maxime Gaudin, Anthony Levasseur, Didier Raoult

**Author notes:** Correspondence to: Pr Didier Raoult, IHU - Méditerranée Infection, 19-21 boulevard Jean Moulin, 13005 Marseille, France. Contributed equally to the work (Philippe Colson, Catherine Dhiver).

## Abstract

Strategies to cure HIV-infected patients by virus-targeting drugs have failed to date. We identified a HIV-1-seropositive woman who spontaneously suppressed HIV replication and had normal CD4-cell counts, no HIV disease, no replication-competent virus and no cell HIV DNA detected with a routine assay. We suspected that dramatic HIV DNA degradation occurred postinfection. We performed multiple nested-PCRs followed by Sanger sequencing and applied a multiplex-PCR approach. Furthermore, we implemented a new technique based on two hybridization steps on beads prior to next-generation sequencing that removed human DNA then retrieved integrated HIV sequences with HIV-specific probes. We assembled ≈45% of the HIV genome and further analyzed the G-to-A mutations putatively generated by cellular APOBEC3 enzymes that can change tryptophan codons into stop codons. We found more G-to-A mutations in the HIV DNA from the woman than in that of her contaminator. Moreover, 74% of the tryptophan codons were changed to stop codons (25%) or were deleted as a possible consequence of gene inactivation. Finally, we found that this woman’s cells remained HIV-susceptible *in vitro*. Our findings show that she does not exhibit innate HIV resistance but has been cured of it by extrinsic factors, a plausible candidate for which is the gut microbiota.

## INTRODUCTION

The evolution of vertebrates has included the integration in their genomes of multiple retrovirus sequences ^1^. Humans are not the exception to the rule as ≈8% of their genome consist in retroviral DNA. Most integrated retrovirus sequences were inactivated by substantial degradation, and only remain as relics of ancient retrovirus epidemics ^2,3^. This general biological phenomenon that consists in the cannibalism of the DNA from viral invaders has been revealed to be on-going in koalas with retroviruses causing an AIDS-like syndrome^4^. Regarding HIV, the problem of prevention and cure of HIV infection has not been solved since its discovery^5,6^. Only two cases of HIV-1 remission have been described following CCR5delta32/delta32 hematopoietic stem-cell transplantation^7,8^. However, some paths could open new therapeutic and preventive avenues. An alternative option to cure patients of HIV might be to strengthen, if identified, natural antiviral defenses. A group of cellular enzymes named APOBEC3 exists whose function is to destroy invading viruses, including retroviruses^9^. The predominant role of APOBEC3s is to deaminate Cs that are changed to Ts, which leads to G-to-A mutations in integrated viral DNA. The genomes of HIV and SIV in great apes encode a protein, Vif, that inhibits their action^10^. However, it has been evidenced in gorillas that a single mutation in the APOBEC3G gene can confer resistance to SIV from chimpanzees by counteracting Vif activity^10^. In addition, it was recently shown that a patient experienced a dramatic decrease in peripheral blood mononuclear cell (PBMC) HIV-1 DNA load in response to a release of immunity by monoclonal antibodies targeting PD-1^11^, whose activity is known to be modulated by the gut microbiota^12,13^. Moreover, the regulation of immune responses by exogenous factors including the microbiota is an emerging field in cancer immunotherapy^12,13^. Thus, the immune control of HIV infection under the influence of exogenous factors is not a theoretical impossibility.

We previously described two HIV-1-seropositive patients who we believe might have spontaneously cured of HIV^14,15^. Indeed, although they never received antiretrovirals, they persistently have a suppressed HIV replication, normal CD4 T cell counts, and no HIV-related disease for more than 10 years (one was HIV-diagnosed in 1985). In addition, no replication-competent HIV was retrieved by culture, and HIV DNA was not detected in peripheral blood mononuclear cells (PBMC) by our diagnosis assay. PBMC HIV DNA was only laboriously obtained by performing hundreds PCR. While searching for other index cases to understand if it is possible to be cured spontaneously of HIV, we investigated a third case.

## METHODS

### Sample collection

Samples were obtained from the patient in January 2015, September 2015 and January 2017 in an attempt to obtain the greatest number of HIV sequences from peripheral blood mononuclear cells (PBMCs). Samples were obtained from the contaminator in January 2017. Informed written consent was obtained from the patients. This study was approved by our institution’s ethics committee (ethics committee of IHU Méditerranée Infection) (N°2018-001).

### PCR amplification, Sanger sequencing and multiplex PCR technique

HIV-1 DNA Sanger population sequencing was performed as described previously^14^. All HIV genes were targeted by at least one PCR system (supplementary information), and all PCRs were conducted in quadruplicate. PCR positivity was determined based on obtaining an HIV sequence by Sanger sequencing. A multiplex PCR technique called “Bortsch” was also performed as described previously^14^.

### Human DNA depletion, HIV-1 DNA enrichment procedures and Illumina next-generation sequencing of DNA extracted from the woman PBMCs

#### Human and HIV-1-specific probe design

Whole human-specific probes (baits) were constructed as described in a previously developed protocol^16^ with modifications^17^. A full-length human genome derived from a modern reference individual (HapMap individual NA21732; Coriell Institute for Medical Research, Camden, NJ) was used as a template to generate biotinylated RNA “bait” libraries spanning the entire human genome. For the design of HIV-1-specific probes, full-length HIV-1 genomes or HIV-1 DNA fragments were fenestrated using a Perl script into a 120 nucleotide-long fragment with a sliding window of 60 nucleotides. The targeted HIV-1 sequences were genomes from the set of HIV-1 reference genomes of the Los Alamos National Institutes of Health HIV sequence database (https://www.hiv.lanl.gov/content/sequence/NEWALIGN/align.html#comp), HIV genomes obtained from two patients whose cases were previously described^14^, and HIV-1 DNA that had been recovered from the woman and her contaminator. The set of 20,000 probes was synthesized by Arbor Biosciences (Arbor Biosciences, Ann Arbor, MI, USA).

#### Library preparation for the next-generation sequencing

DNA extraction was performed on 200 µL of PBMCs (≈2e5 cells) collected from the woman, using the EZ1 Virus Mini Kit v2.0 (Qiagen Hilden, Germany) according to the manufacturer’s protocols. Five paired-end libraries were prepared using 1 ng of extracted DNA and MiSeq Technology with the paired-end method and the Nextera XT kit (Illumina Inc., San Diego, CA, USA). DNA was fragmented, and adaptors containing the Illumina P5/P7 primer sequences and tags were added.

#### Human DNA depletion procedure

Five depletions of human nucleic acids were performed separately by hybridization of 50 µl of each prepared Illumina library (≈100 ng of DNA) with 500 ng of biotinylated human RNA-bait library. Targeted fragment/probe heteroduplexes were captured using magnetic streptavidin-harboring beads (MyOne Streptavidin C1 Dynabeads (Life Technologies, Carlsbad, USA)), as previously described^17^. The unbound fraction (supernatant) was concentrated and cleaned using 1.8× AMPure XP beads (Beckman Coulter, Fullerton, CA, USA) according to the manufacturer’s protocol with elution into 30 µl of 1X TE buffer. The five purified fractions were then pooled and concentrated using a MinElute PCR Purification Kit (Qiagen) according to the manufacturer’s protocol with elution in 10 µl of elution buffer. To generate sufficient material for targeted enrichment, this product was amplified using eight PCR amplification cycles with Illumina P5/P7 primers, before purification using a MinElute PCR Purification Kit (Qiagen) and elution with 10 µl of elution buffer.

#### Targeted HIV enrichment through hybridization capture

A total of 500 ng of the human-depleted library was used to perform the targeted HIV enrichment step involving hybridization with the HIV-specific probes using a myBaits target capture kit (Arbor Biosciences) according to the manufacturer’s instructions (Hybridization Capture for Targeted NGS manual version 4.01). Hybridization-based capture reactions with undiluted HIV-1 probes (500 ng) was performed at 65°C for 16 h. Streptavidin-coated magnetic beads (myBaits kit) were added to the hybridization mixture, and the sample was additionally incubated for 5 min at 65°C. After washing steps, beads were resuspended in 30 µL of 10 mM Tris-Cl, 0.05% Tween-20 solution (pH 8.0-8.5). The captured DNA was dissociated from beads by heating the suspension at 95°C for 5 min. HIV-1 DNA and human albumin DNA were quantified by a multiplex real-time PCR assay as previously described^14^.

#### Next-generation sequencing and sequence read analysis

The product of the HIV enrichment procedure was normalized according to the Nextera XT protocol for pooling and sequencing on a MiSeq instrument (Illumina). A single run of 39 h in 2×250 base pairs (bp) was carried out for paired-end sequencing and cluster generation. Reads were filtered based on their quality, generated paired reads were imported into the CLC software (https://www.qiagenbioinformatics.com/products/clc-genomics-workbench/)and then assembled by mapping to the HIV genome GenBank accession no. K03455.1 (HIV-1 strain HXB2) and to sequences obtained from the woman and contaminator PBMCs. Reads identified as corresponding to HIV sequences were exported as fasta and SAM files.

### Comparisons of HIV-1 sequences obtained from the woman PBMCs and her contaminator PBMCs and analysis of G-to-A mutations and of substitution of tryptophan codons by stop codons

A custom script written in Python language was used to analyze the SAM file generated from the mapping of reads obtained by next-generation sequencing and to count differences in amino acids between HIV sequences recovered from the PBMCs of the woman and her contaminator. In addition, alignments of nucleotide sequences obtained by Sanger and next-generation sequencing were performed using the MUSCLE program^18^. The phylogenetic analysis was performed using the MEGA6 software (www.megasoftware.net) with the neighbor-joining method. Nucleotide sequences were merged into a Microsoft Excel file using sequences from the HIV-1 K03455.1 genome (HXB2 strain) or those obtained from the contaminator PBMCs as references. An alignment was also performed using the same sequences after their translation into amino acids from the three open reading frames using the Transeq online tool (https://www.ebi.ac.uk/Tools/st/emboss_transeq/) and amino acid sequences from the proteins of the HXB2 strain as references, and these aligned sequences were merged into the Microsoft Excel file. G-to-A mutations and substitution of tryptophan codons by stop codons were searched using the Microsoft Excel software.

### HIV culture assay

Testing for PBMC resistance to HIV was conducted as previously described^14,19^.

### Supplementary material

HIV sequences obtained in the present study are available at https://www.mediterranee-infection.com/acces-ressources/donnees-pour-articles/hiv/ or have been submitted to GenBank (submission ID: 2225641).

## RESULTS

The patient is a 37-year-old woman sexually infected with HIV-1 between 2002 and 2004 by a single partner and diagnosed as seropositive in July 2006 (see Supplementary Fig. S1 online). She never received antiretrovirals except in 2012 during the third trimester of her pregnancy (zidovudine, 300 mg/d). Nevertheless, to date, she has persistently had normal CD4 T lymphocyte counts (mean value between 2006 and 2018, 1,221±172/mm^3^) and remained free of HIV-related disease. In addition, HIV RNA was not detected in plasma using commercialized PCR assays on ten occasions during follow-up, and no replication-competent HIV was retrieved by coculture. Moreover, no PBMC HIV DNA was detected by routine diagnostic tests^14^ on six occasions between 2010 and 2018. However, the woman’s PBMCs were found to be susceptible to the HIV-1 NL4-3 strain. This finding indicates that this woman acquired the capability to combat HIV after her infection and suggests the role of an extrinsic factor. Moreover, she was not infected with a defective HIV strain, as in her contaminator, a 30-year-old man HIV-diagnosed in 1990, the HIV DNA load was 350 copies/million PBMCs, and the CD4 T cell count fell to <200/mm^3^, which required antiretroviral therapy.

Based on these findings and our previous work^14^, we suspected that the HIV genome in this young woman had been drastically degraded after its integration. We attempted to obtain fragments of HIV DNA from her PBMCs and assess their degradation by using thorough molecular procedures and the contaminator’s HIV sequences as a reference. We used different strategies to retrieve the maximum number of HIV sequences from the PBMCs of this woman in whom standard assays had failed to detect HIV DNA. First, we performed nested PCR targeting HIV sequences from the literature, including with a multiplex PCR technique^14^. During these steps, HIV sequences were obtained by Sanger sequencing from 26 (7%) of 392 nested PCRs performed on the woman PBMCs (Table 1). The mean PCR product size was 251±201 nucleotides, 54% being shorter than 200 nucleotides, and they assembled into 3,254 nucleotide-long noncontiguous fragments classified as HIV-1 subtype B (Fig. 1). HIV sequences obtained from the woman matched those of the near full-length HIV genome (9,337 nucleotides) recovered from the PBMCs of her partner, confirming that this patient was the contaminator and that the young woman was infected with a nondefective viral strain and eliminating a contamination. Second, for the first time, we implemented a technique using two successive steps of uptake on probe-coated beads. The two steps consisted of human DNA depletion with human DNA-targeting probes followed by HIV DNA enrichment with a set of HIV probes complementary to the contaminator HIV sequences. This enriched HIV DNA was thereafter nonspecifically amplified and sequenced by Illumina next-generation sequencing. We obtained 73 reads through this procedure that increased the length of the assembled HIV genome by 17% (1,133 nucleotides) (Fig. 2a; see Supplementary Information online). Overall, by these three approaches and carrying out hundreds of manipulations, we obtained a set of noncontiguous fragments covering 4,387 nucleotides of the integrated HIV DNA of this young woman.

**Table 1.**
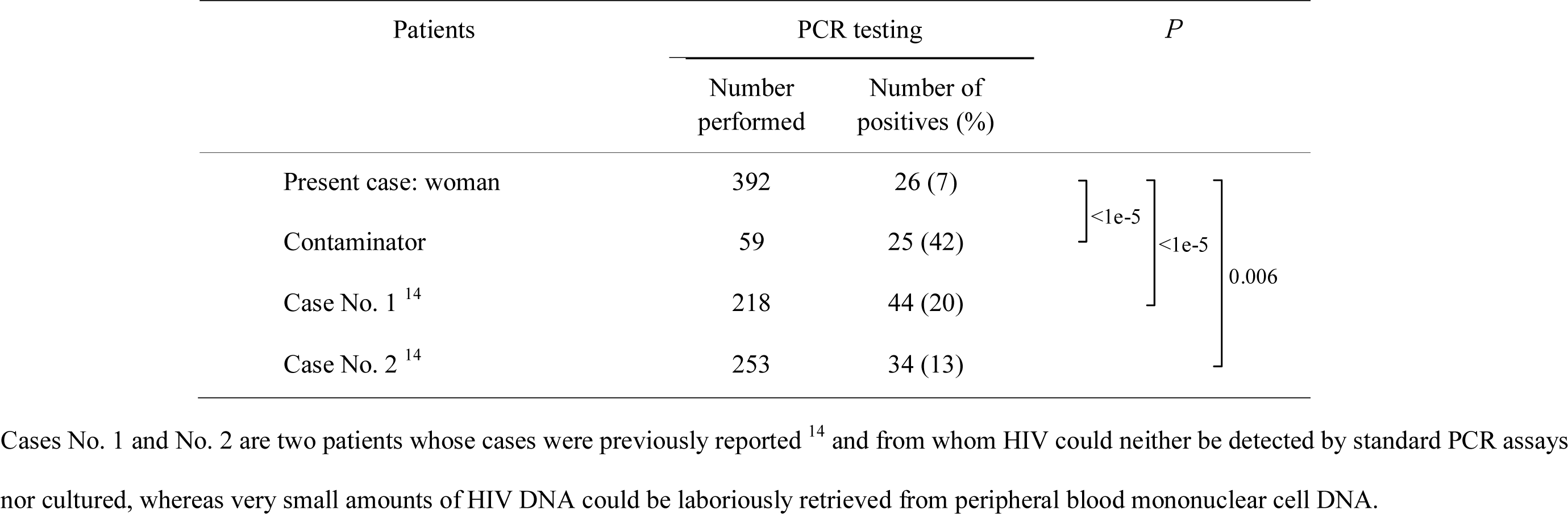
Proportions of positive HIV DNA test results for PCR systems used on DNA of peripheral blood mononuclear cells from HIV seropositive cases and controls

**Figure 1.**
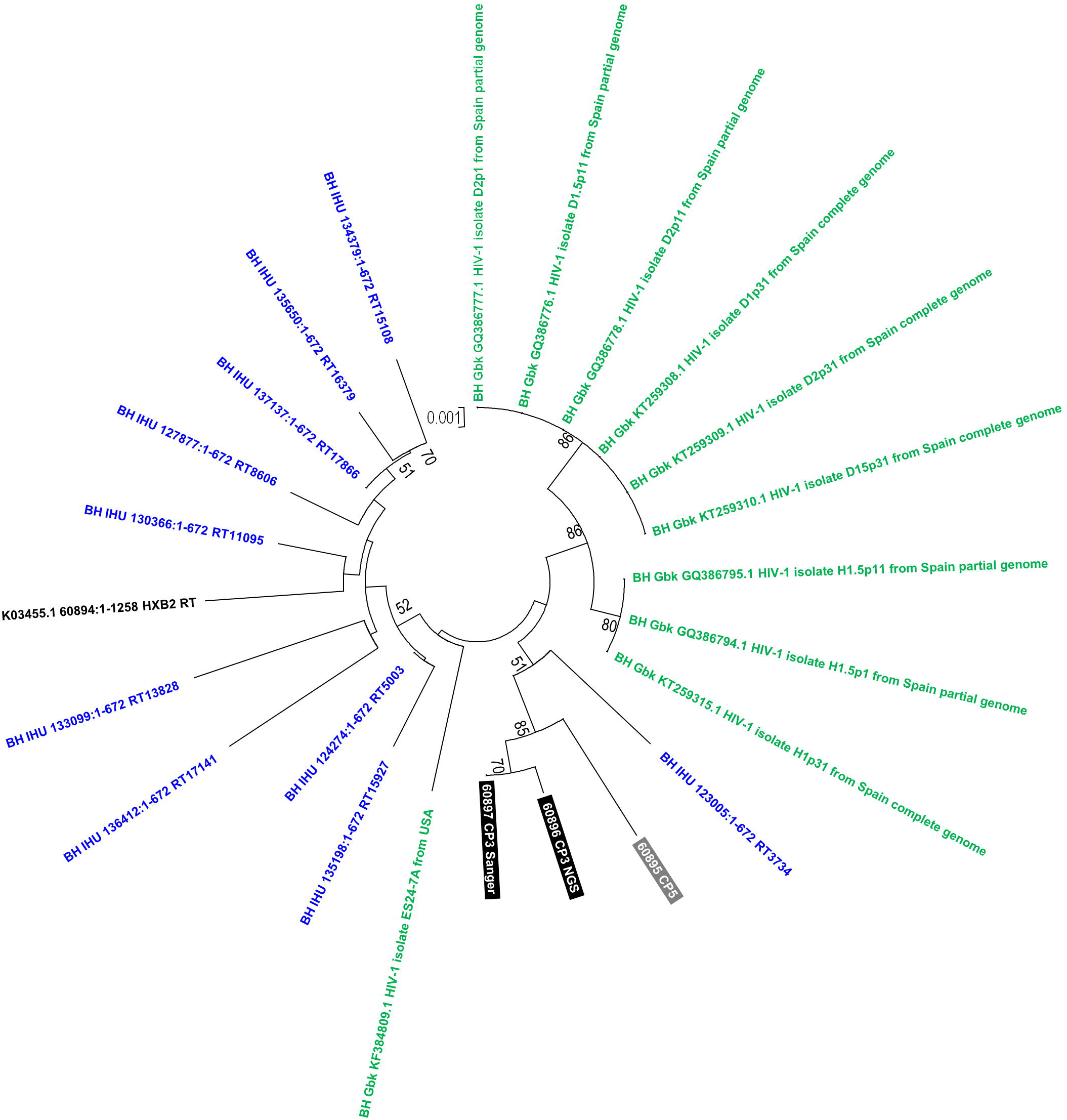
Phylogenetic analysis of HIV reverse transcriptase sequences obtained from the PBMC DNA of the woman and her contaminator. The HIV-1 genome fragment analyzed here corresponds to a 671-nucleotide alignment generated from sequences of the reverse transcriptase-encoding gene and corresponding to nucleotides 2,596-3,266 of the HIV-1 genome GenBank accession no. K03455.1. Sequences obtained from the present cases are indicated by a bold white font and a black (CP3, woman) or a gray (CP5, contaminator) background. The 10 sequences with the highest BLAST score recovered from the NCBI GenBank nucleotide sequence database (http://www.ncbi.nlm.nih.gov/nucleotide/), labeled with BH Gbk (for best BLAST hit GenBank) and indicated by a green font, and from our local sequence database, labeled with BH IHU (for best BLAST hit IHU-Méditerranée Infection) and indicated by a blue font, were incorporated in the phylogeny reconstruction. Nucleotide alignments were performed using the MUSCLE software (http://www.ebi.ac.uk/Tools/msa/muscle/). The evolutionary history was inferred in the MEGA6 software (http://www.megasoftware.net/) using the neighbor-joining method and the Kimura 2-parameter method. The percentage of replicate trees in which the associated taxa clustered together in the bootstrap test (1,000 replicates) is shown next to the branches. The tree is drawn to scale, with branch lengths in the same units as those of the evolutionary distances used to infer the phylogenetic tree; the scale bars indicate the number of nucleotide substitutions per site. Bootstrap values >50% are labeled on the tree. NGS, next-generation sequencing.

**Figure 2.**
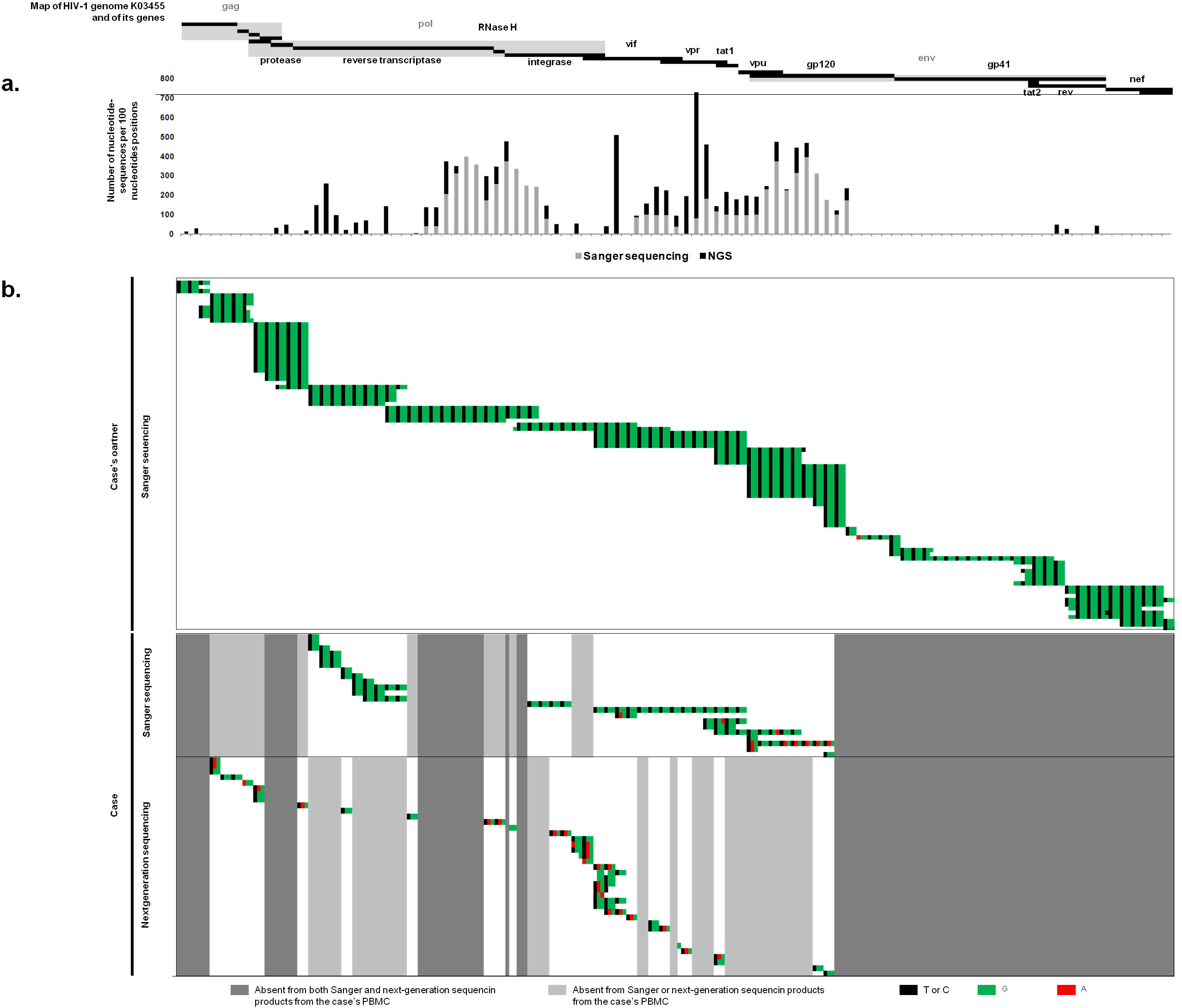
Condensed view of the location and number of HIV DNA fragments recovered from the PBMCs of the woman using Sanger and next-generation sequencing (a) and coverage of tryptophan stop codons in the HIV-1 genome by HIV sequences retrieved from the woman and contaminator PBMCs by Sanger or next-generation sequencing (b). a: The number of nucleotide sequences per 100 nucleotide positions corresponds to the sum of sequences covering each nucleotide position per window of 100 nucleotides. b: All 91 tryptophan stop codons covered by Sanger sequencing products obtained from the contaminator’s PBMCs. Green indicates G nucleotides, red indicates A nucleotides, black indicates T or C nucleotides. Regarding sequences generated from the woman PBMCs, areas in dark and light gray indicate tryptophan codons not covered by any sequences obtained by both Sanger and next-generation sequencing or by one of these sequencing strategies, respectively.

Then, as the genome of the HIV strain that infected this young woman had been obtained from her contaminator, we were able to determine the number of G-to-A mutations attributed to APOBEC3G activity in her HIV DNA, and specifically the number of tryptophan codons eliminated either by their change to stop codons or because fragments of DNA containing the tryptophan codons were lost. G-to-A mutations were observed in the woman’s HIV DNA at 152 (16%) positions of the HIV DNA from the contaminator that only harbored G, and a G-to-A excess was detected in the woman compared to her contaminator, notably for the genes encoding HIV reverse transcriptase; integrase; Vif, which counteracts APOBEC3G; Vpr; Vpu; and Env gp120 (Table 2; see see Supplementary Table S1 online). We thereafter determined that HIV sequences of the woman only covered 47 (52%) of the 91 tryptophan codons of the HIV genome; however, they were all present in sequences retrieved from the contaminator (Table 3; Fig. 2b). In the woman’s HIV DNA, changes from a tryptophan codon to a stop codon were observed 5 times in *vif* and *gp120* envelope genes, 4 times in integrase gene, and twice in *gag*, reverse transcriptase, *vpr* and *vpu* genes, contrasting with no such changes in HIV DNA from the contaminator. With regard to HIV sequences generated by next-generation sequencing of DNA from the woman PBMCs, 67 (74%) of the 91 tryptophan codons present in the contaminator HIV DNA were either retrieved but changed to stop codons (as observed for 17 (63%) of 27 tryptophan codons on average (Table 4)) or located in HIV-1 DNA regions that we did not retrieve as a possible consequence of their degradation or loss. Thus, G-to-A mutations, the absence of codon coverage, or both states combined were significantly more frequent at tryptophan codons than at any other codon. Overall, we found that G-to-A mutations generating stop codons occurred in at least one sequence at 23 (49%) of the 47 covered tryptophan codons (at 9 and 16 of those covered by Sanger and next-generation sequencing, respectively) in viral sequences recovered from the woman, whereas no such change was detected in HIV sequences obtained from the contaminator at any of the 91 covered tryptophan codons. Taken together, these findings suggest that APOBEC3G activity was greater in the woman than in her contaminator and this increased activity induced a dramatic degradation of integrated PBMC HIV DNA after infection. The detection of several stop codons in the Vif-encoding gene is particularly worthy of note because this protein counteracts APOBEC3G by triggering its degradation^20^. In addition, APOBEC3G DNA sequencing in the patient did not show mutations compared to reference sequences at Vif-APOBEC3 interaction sites, oligomerization/encapsidation sites, and N- and C-terminal active sites (see see Supplementary Fig. S2 online).

**Table 2.** G-to-A mutations detected in HIV DNA fragments obtained by Sanger or next-generation sequencing from woman PBMCs at G-harboring positions in the contaminator’s HIV DNA

| HIV-1 gene | Number of positions<br>harboring G-to-A mutations |  |
| --- | --- | --- |
|  | Number | % |
| *gag* | 15 | 12 |
| *pol protease* | 6 | 15 |
| *pol reverse transcriptase* | 20 | 9 |
| *pol RNase H* | 6 | 29 |
| *pol integrase* | 45 | 26 |
| *vif* | 30 | 23 |
| *vpr* | 13 | 17 |
| *tat* | 4 | 9 |
| *vpu* | 17 | 33 |
| *rev* | 4 | 11 |
| *env gp120* | 17 | 19 |
| *env gp41* | 1 | 3 |
| *nef* | 2 | 13 |

**Table 3.** Tryptophan-to-stop codon mutations detected in HIV DNA fragments obtained from the woman and contaminator PBMCs by Sanger and next-generation sequencing

| HIV-1 gene | Number of sequences harboring mutations<br>generating tryptophan codon-to-stop codon<br>changes |  |
| --- | --- | --- |
|  | Woman PBMCs<br>(Sanger/NGS) | Contaminator PBMCs |
| *gag* | -/2 | 0 |
| *pol protease* | -/1 | 0 |
| *pol reverse transcriptase* | 0/2 | 0 |
| *pol integrase* | 0/4 | 0 |
| *vif* | 1/5 (1 in common) | 0 |
| *vpr* | 1/2 (1 in common) | 0 |
| *tat* | 0/- | 0 |
| *vpu* | 2/0 | 0 |
| *rev* | -/- | 0 |
| *env gp120* | 5/0 | 0 |
| *env gp41* | -/- | 0 |
| *nef* | -/- | 0 |

-, no sequence; NGS, next-generation sequencing

**Table 4.** G-to-A mutations detected in HIV DNA fragments obtained from the woman PBMCs by next-generation sequencing in reference to HIV DNA obtained from the contaminator PBMCs

| HIV-1 codons * | Numbers |  |  |  | P |  |  |  |
| --- | --- | --- | --- | --- | --- | --- | --- | --- |
|  | Total in the HIV genome | Covered by NGS reads | Not mutated (mean) | Mutated (mean) | Uncovered vs covered by NGS reads | Mutated vs not mutated | Lost vs not mutated | Mutated or lost vs not mutated |
| A | 188 | 59 | 36 | 23 | - | - | - | - |
| C | 55 | 14 | 12 | 2 | - | - | - | - |
| D | 125 | 35 | 29 | 6 | - | - | 0.0388 | - |
| E | 227 | 75 | 45 | 30 | - | - | - | - |
| F | 86 | 26 | 19 | 7 | - | - | - | - |
| G | 226 | 80 | 57 | 23 | - | - | - | - |
| H | 72 | 22 | 16 | 7 | - | - | - | - |
| I | 212 | 77 | 48 | 29 | - | - | - | - |
| K | 211 | 77 | 56 | 21 | - | - | - | - |
| L | 260 | 74 | 55 | 20 | - | - | - | - |
| M | 62 | 24 | 15 | 9 | - | - | - | - |
| N | 147 | 42 | 34 | 8 | - | - | - | - |
| P | 176 | 62 | 41 | 21 | - | - | - | - |
| Q | 180 | 65 | 45 | 20 | - | - | - | - |
| R | 192 | 52 | 33 | 19 | - | - | - | - |
| S | 168 | 56 | 39 | 17 | - | - | - | - |
| T | 170 | 46 | 36 | 10 | - | - | - | - |
| V | 184 | 65 | 40 | 25 | - | - | - | - |
| **W** | **88** | **27** | **10** | **17** | - | **0.0189** | **0.0009** | **0.0051** |
| Y | 80 | 33 | 23 | 10 | - | - | - | - |

-, >0.05; \* Named according to the IUPAC amino acid notation (https://iupac.org/; W= tryptophan); NGS, next-generation sequencing

## DISCUSSION

We present here evidence that a young woman was infected with HIV, as she is seropositive and HIV sequences were eventually retrieved from her PBMCs using ultrasensitive methods, but after infection spontaneously and thoroughly degraded the HIV genomes integrated in her DNA. This occurred through G-to-A mutations, which is the signature of APOBEC3 cellular enzymes, and led to gene inactivation by changing tryptophan codons into stop codons. Clinically, the woman never developed immunodeficiency or HIV-related symptoms, which suggests that she was cured of HIV^15^. The fate of HIV infection was totally different in her contaminator although both individuals where infected with a same HIV strain. These findings highlight that the different outcome relied on host response to infection, not on the viral strain. The woman harbored far less abundant and more degraded HIV DNA than her contaminator, revealing a more extensive action of APOBEC3 enzymes. In humans, greater APOBEC3G amounts in blood resting memory CD4 cells was associated with lower PBMC HIV DNA levels^21^. In addition, increased rates of G-to-A mutations were observed among HIV-seropositive individuals who spontaneously suppress HIV replication^14,22^.

The present case is critical because it shows that some individuals are likely to inactivate integrated HIV after infection. As this woman’s cells remained susceptible to HIV infection, this phenomenon appears to be linked to extrinsic factors. A possible explanation is that the microbiota of this woman hyperactivated APOBEC3 enzymes. Preliminary work has highlighted that bacterial components could stimulate the expression of APOBEC3G and enhance its activity on integrated HIV, which is certainly a path worthy of exploration^23^. The modulation of the immune response by the digestive microbiota, in particular in the ileum where lymphocytes pass several times per day, is one of the keys that may open the way to new therapeutic strategies to fight HIV, as has been reported in the oncology field for which specific gut microbes are shown to drastically modulate responses to cancer immunotherapies.^13^.

## Contributors

DR and PC designed the study. CD provided clinical data. PC, JD, OOG, MG and AL performed the molecular biology or bioinformatic analyses. PC, CD, CT, AL and DR analysed the study data. DR and PC wrote the manuscript. All authors revised or reviewed the manuscript critically and approved the final version.

## Funding

This work was supported by the French Government under the “Investments for the Future” program managed by the National Agency for Research (ANR), Méditerranée-Infection 10-IAHU-03 and was also supported by Région Provence Alpes Côte d’Azur and European funding FEDER PRIMMI (Fonds Européen de Développement Régional - Plateformes de Recherche et d’Innovation Mutualisées Méditerranée Infection).

## Role of the funding source

The funders of the study had no role in study design, data collection, data analysis, data interpretation, and writing of the report. All authors had full access to the data in the study and the corresponding author had final responsibility for the decision to submit for publication.

## Competing interests

The authors declare no competing interests.

## Acknowledgments

We thank Annick Abeille, Emilie Doudon, Emeline Baptiste, and Nathalie Duclos for their technical assistance.

