## Supplementary Information for "Dramatic HIV DNA degradation associated with spontaneous HIV suppression and disease-free outcome in a young seropositive woman following her infection"

**SUPPLEMENTARY Methods**

**PCR amplification and sequencing**

HIV-1 DNA was extracted from peripheral blood mononuclear cells (PBMCs) of the two individuals using the EZ1 viral nucleic acid extraction kit (DNA tissue kit, Qiagen, Hilden, Germany) according to the manufacturer’s recommendations, following a preliminary step with digestion by proteinase K at 56°C for 60 min, as previously described^1^. Nested PCR amplification was performed using FastStart Taq DNA polymerase (Roche Applied Science, Indianapolis, IN, USA), 0.4 mM of each dNTP, 3.2 mM MgSO_4_ and 200 nM of each primer. The cycling conditions were as follows: 94°C for 4 min, then 39 cycles of 30 sec at 95°C, 45 sec at the most appropriate temperature according to primers used and 2 min at 72°C, before a final elongation step at 72°C for 10 min. Primers used are listed in table S2. DNA from approximately 200,000 PBMCs was tested per PCR. HIV-1 DNA sequences were obtained via Sanger population sequencing using the Big Dye Terminator cycle sequencing kit version 1.1 on the ABI Prism 3130 xl genetic analyzer (Applied Biosystems, Branchburg, NJ, USA) under the conditions previously described^1^ and were analyzed using SeqScape v2.5 (Applied-Biosystems) or ChromasPro (http://technelysium.com.au/?page_id=27) softwares. Based on the alignment of the HIV-1 genomes that were the best BLAST hits of HIV sequences obtained from the two individuals’ PBMCs, conserved regions were identiﬁed and used as targets to design PCR primers. HIV-1 DNA next-generation sequencing using the “Bortsch” procedure was performed as previously described^1^ by performing PCR amplification in a single tube reaction with a mix of several primers with the aim of obtaining a “pool” of fragments of different lengths and then submitted to next-generation sequencing.

**Analysis of sequences obtained by Sanger sequencing**

To obtain a consensus sequence from HIV DNA fragments retrieved from the patient and contaminator PBMCs, these sequences were mapped to the HIV genome GenBank Accession no. K03455 using the CLC Bio software (www.clcbio.com/). Then, an alignment was performed using the MUSCLE tool from the MEGA v6.0 platform^2^ that was manually curated before consensus sequence generation using the SeaView program^3^.

### Human DNA depletion and HIV-1 DNA enrichment for DNA extracted from woman PBMCs

### Real-time PCR targeting the human beta-actin gene was performed on the DNA extracted from the woman PBMC, as published previously^4^.

### Analysis of sequence reads generated by next-generation sequencing

### Mapping of sequence reads generated by next-generation sequencing to the HIV genome assembled from viral sequences retrieved from the PBMCs of the contaminator was performed using CLC Bio software 7.5 (http://www.clcbio.com) with the following parameters: similarity fraction 0.7 and length fraction 0.9. The CLC mapping report was then used to analyze single nucleotide polymorphisms between mapped reads and the reference. Two amino acids included in the frameshift in *tat* and *rev* genes were ignored because no read covered them.

**HIV culture assay**

Testing for PBMC resistance to HIV was conducted as previously described^1, 5^. Briefly, 2 x 10^6^ PHA-prestimulated patient PBMCs suspended in 2 ml of RPMI-1640 medium with 20% fetal calf serum, 0.4 IU/ml interleukin-2, L-glutamine and penicillin were used. The HIV NL4.3 strain was used for inoculation. The cultures were maintained for 10 days, and cell-free supernatants were harvested twice a week and tested by real-time PCR (generic HIV RNA assay). PBMCs not inoculated with HIV were used as negative control.

**Sequencing of the gene encoding APOBEC3G**

Sequencing of the gene encoding APOBEC3G was performed from the woman PBMCs. DNA extraction was performed using the EZ1 Virus Mini Kit v2.0 (Qiagen) according to the manufacturer’s protocols. The PCR primers used are indicated in table S3. Sequences were obtained via Sanger population sequencing using the Big Dye Terminator cycle sequencing kit version 1.1 on the ABI Prism 3130 xl genetic analyzer (Applied Biosystems, Branchburg, NJ, USA), and were analyzed using ChromasPro (http://technelysium.com.au/?page_id=27) software.

**Sequencing of the C-C chemokine receptor 5 (CCR5)**

Sequencing of the gene encoding the C-C chemokine receptor 5 (CCR5) was performed from the case-patient PBMCs to search for the 32-base pair deletion using previously described PCR primers^6^.

**Statistical analyses**

Statistical analyses were performed with the Openepi software (v3.03a; http://www.openepi.com). Proportions were compared using the chi-square test or the Fisher test. P< 0.05 was considered significant.

**SUPPLEMENTARY RESULTS**

### Human DNA depletion, HIV-1 DNA enrichment, Illumina next-generation sequencing of DNA extracted from woman PBMCs, and sequence analysis

### Real-time PCR targeting the actin gene that was performed on the DNA extracted from the woman PBMCs was positive with a cycle threshold (Ct) of 23, whereas no HIV-1 DNA was detected using our routine real-time qPCR diagnostic assay^1^. After Illumina library preparation and human DNA depletion, the actin gene was no longer detected. Illumina sequencing performed following the HIV-1 DNA enrichment step generated a total of 146 reads that were mapped to the HIV genome assembled from viral sequences retrieved from the PBMCs of the contaminator. As forward and reverse reads mapped to the same coordinates, 73 unique reads corresponding to HIV sequences were kept for further analyses. No read matching the HIV genome was found in a “blank” control sample with no DNA input.

### HIV sequences availability

### HIV sequences obtained from the two patients are available at the following URL: https://www.mediterranee-infection.com/acces-ressources/donnees-pour-articles/hiv/ (file: "5files") and have been submitted to the NCBI GenBank nucleotide sequence database (submission ID: 2225641 (CP3= woman; CP5= contaminator)).

**HIV culture assay**

HIV culture assays showed that the PBMCs from the woman were susceptible to infection by the HIV-1 NL4.3 strain (Fig. S3).

**Sequencing of the gene encoding C-C chemokine receptor 5 (CCR5)**

Sequencing of the gene encoding CCR5 showed sequences devoid of the 32-base pair deletion.

**Supplementary Figures**

**Figure S1.** HIV-1 western blot testing in the HIV-1-seropositive woman.

HIV-1 western blot testing was performed on a serum sample collected in January 2015 using the Bio-Rad kit (Stanford, CA, USA), following the manufacturer’s recommendations. Negative and positive controls were samples from the kit.

bp, binding protein; em, extramembranous; Env, envelope; Int, integrase; kDa, kilodalton; MA, matrix; MW, molecular weight; NC, nucleocapsid; pre, precursor; Prot, protease; RT, reverse transcriptase; su; subunit; Tev, fusion of Tat, Env and Rev; tm, transmembranous

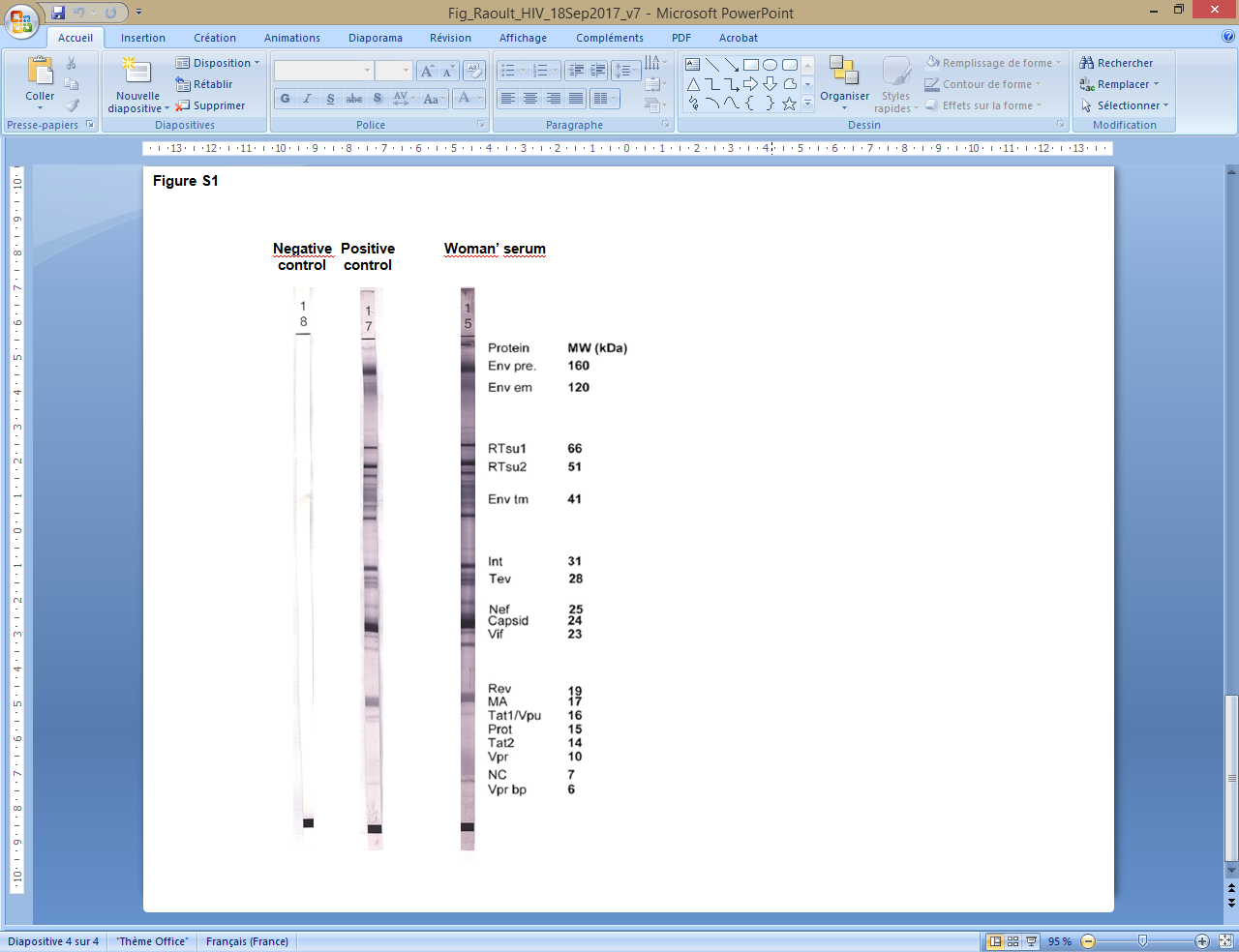

**Figure S2.** Sequences of the human APOBEC3G-encoding gene in regions previously identified as having a functional role, including interaction with HIV Vif protein^7^.

The GenBank accession number NM_021822.3 sequence was used as a reference.

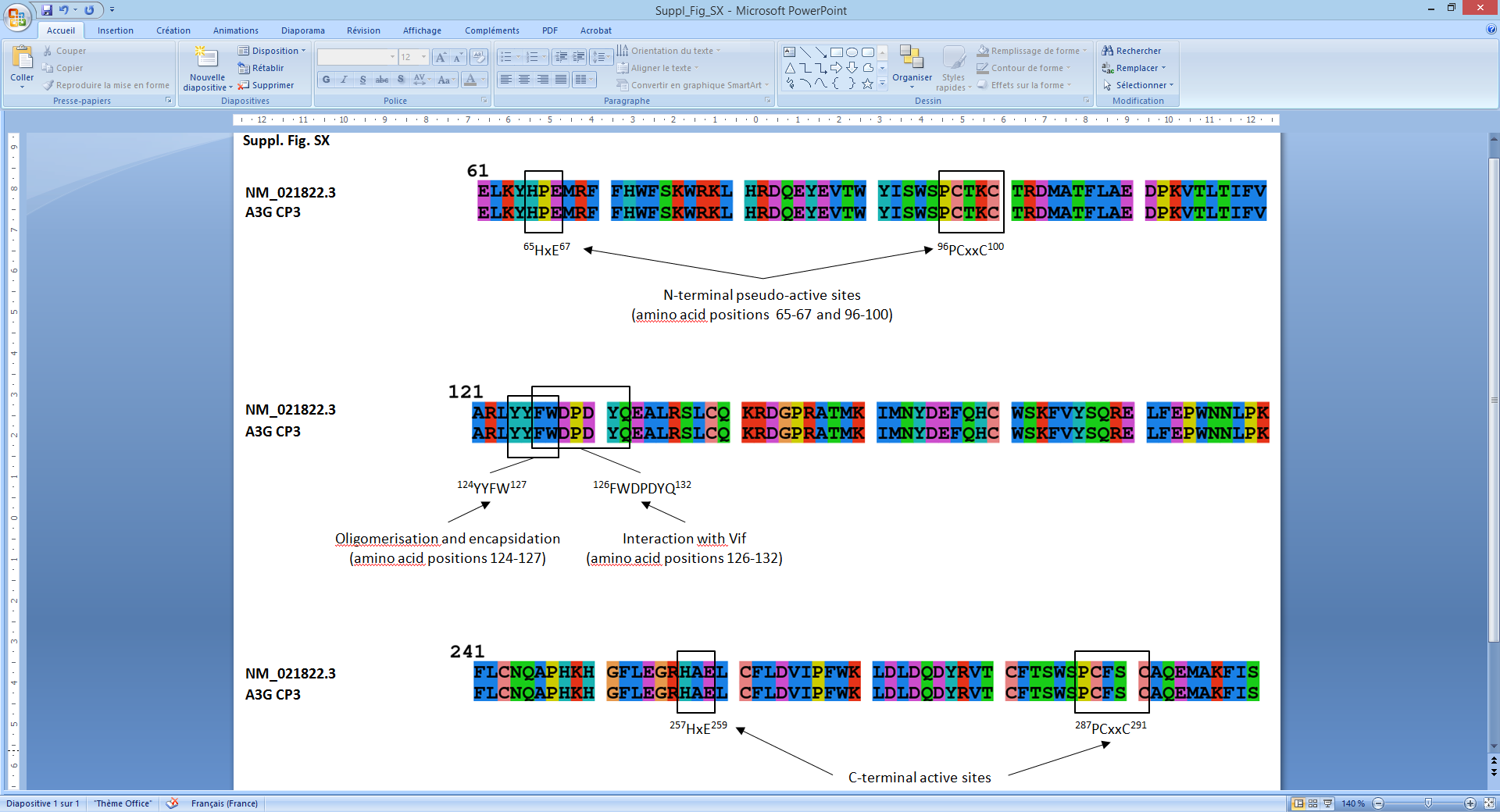

**Figure S3.** HIV-1 DNA (a) and RNA (b) assessment of susceptibility to infection by the HIV-1 NL4.3 strain of PBMCs from the woman by HIV culture assays

a.

HIV-1 DNA (log10 copies/106 PBMC)

| Days post-HIV-1 inoculation | Negative control | Test #1 | Test #2 |
| --- | --- | --- | --- |
| 3 | 0 | 3.79934 | 2.75511 |
| 4 | 0 | 2.41497 | 2.5611 |
| 5 | 0 | 3.2878 | 3.29003 |
| 6 | 0 | 2.37658 | 2.4456 |
| 7 | 0 | 3.15229 | 3.07188 |

b.

HIV-1 RNA (log10 copies/mL)

| Days post-HIV-1 inoculation | Negative control | Test #1 | Test #2 |
| --- | --- | --- | --- |
| 3 | 0 | 6.59329 | 6.1038 |
| 4 | 0 | 6.34635 | 6.04532 |
| 5 | 0 | 7.17319 | 7.05308 |
| 6 | 0 | 7.99034 | 7.59329 |
| 7 | 0 | 8.63448 | 8.51851 |

**Supplementary Tables**

**Table S1.** G-to-A mutations detected in HIV DNA fragments obtained by Sanger sequencing from woman PBMCs and contaminator PBMCs at G-harboring positions in the HXB2 strain genome

| HIV-1 gene |  | Number of G-harboring positions in the HXB2 strain genome |  | Number (%) of positions harboring A as majority nucleotide in HIV DNA fragments | | P |
| --- | --- | --- | --- | --- | --- | --- |
|  |  |  |  | Woman PBMCs | Contaminator PBMCs |  |
| *pol protease* |  | 22 |  | 0 (0) | 1 (5) | 0.5 |
| *pol reverse transcriptase* |  | 210 |  | 44 (21) | 11 (5) | <10^-3^ |
| *pol integrase* |  | 136 |  | 46 (34) | 5 (4) | <10^-3^ |
| *vif* |  | 139 |  | 54 (39) | 10 (7) | <10^-3^ |
| *vpr* |  | 86 |  | 35 (41) | 7 (8) | <10^-3^ |
| *tat* |  | 47 |  | 14 (30) | 2 (4) | <10^-3^ |
| *vpu* |  | 58 |  | 21 (36) | 4 (7) | <10^-3^ |
| *rev* |  | 19 |  | 8 (42) | 1 (5) | 0.02 |
| *env gp120* |  | 80 |  | 29 (36) | 8 (10) | <10^-3^ |

GenBank accession no. for the HXB2 strain HIV-1 genome is K03455.1

**Table S2.** PCR primers used to amplify and sequence HIV-1 sequences from the PBMCs of the woman and the contaminator

| Name | Orientation | Best hit in GenBank | | |  | Coordinates in HXB2 strain (K03455.1) | |
| --- | --- | --- | --- | --- | --- | --- | --- |
|  |  | Identification | Start coordinate | End coordinate |  | Start | End |
| HIV_COH__1 | Reverse | gi\|874510288\|gb\|KP877266.1\| | 351 | 334 |  | 2603 | 2586 |
| HIV_COH__2 | Reverse | gi\|887517014\|gb\|KT074935.1\| | 1760 | 1741 |  | 2578 | 2559 |
| HIV_COH__3 | Forward | gi\|870833146\|gb\|KP178444.1\| | 1349 | 1366 |  | 2171 | 2188 |
| HIV_COH__4 | Forward | gi\|870833116\|gb\|KP178443.1\| | 1437 | 1456 |  | 2211 | 2230 |
| HIV_COH__5 | Forward | gi\|874510300\|gb\|KP877272.1\| | 293 | 312 |  | 2545 | 2564 |
| HIV_COH__6 | Forward | gi\|822094503\|gb\|KP455640.1\| | 5020 | 5039 |  | 5862 | 5881 |
| HIV_COH__7 | Reverse | gi\|822094503\|gb\|KP455640.1\| | 5039 | 5020 |  | 5881 | 5862 |
| HIV_COH__8 | Forward | gi\|869135537\|gb\|KR822835.1\| | 5390 | 5409 |  | 5956 | 5975 |
| HIV_COH__9 | Forward | gi\|194031667\|gb\|EU602876.1\| | 543 | 555 |  | 2913 | 2921 |
| HIV_COH__10 | Reverse | gi\|194031667\|gb\|EU602876.1\| | 555 | 543 |  | 2921 | 2913 |
| HIV_COH__11 | Reverse | gi\|675118547\|gb\|KJ888885.1\| | 194 | 172 |  | 1261 | 1240 |
| HIV_COH__12 | Forward | gi\|887517014\|gb\|KT074935.1\| | 2875 | 2894 |  | 3693 | 3712 |
| HIV_COH__13 | Reverse | gi\|877797356\|emb\|LM654312.1\| | 341 | 322 |  | 4558 | 4539 |
| HIV_COH__14 | Reverse | gi\|887517014\|gb\|KT074935.1\| | 3382 | 3357 |  | 4200 | 4175 |
| HIV_COH__15 | Forward | gi\|871044527\|gb\|KP873164.1\| | 3282 | 3301 |  | 3771 | 3787 |
| HIV_COH__16 | Reverse | gi\|887517014\|gb\|KT074935.1\| | 3655 | 3636 |  | 4473 | 4454 |
| HIV_COH__17 | Reverse | gi\|887517014\|gb\|KT074935.1\| | 3362 | 3343 |  | 4180 | 4161 |
| HIV_COH__18 | Forward | gi\|869135503\|gb\|KR822832.1\| | 4916 | 4932 |  | 5520 | 5536 |
| HIV_COH__19 | Forward | gi\|869135478\|gb\|KR822830.1\| | 4796 | 4817 |  | 5438 | 5459 |
| HIV_COH__20 | Forward | gi\|871044527\|gb\|KP873164.1\| | 5077 | 5095 |  | 5563 | 5581 |
| HIV_COH__21 | Reverse | gi\|883744188\|gb\|KT008660.1\| | 246 | 227 |  | 6473 | 6454 |
| HIV_COH__22 | Reverse | gi\|658309113\|emb\|HG327288.1\| | 182 | 161 |  | 6394 | 6382 |
| HIV_COH__23 | Reverse | gi\|887517014\|gb\|KT074935.1\| | 5535 | 5514 |  | 6357 | 6336 |
| HIV_COH__24 | Forward | gi\|887517014\|gb\|KT074935.1\| | 5520 | 5538 |  | 6342 | 6360 |
| HIV_COH__25 | Forward | gi\|871044527\|gb\|KP873164.1\| | 6038 | 6058 |  | 6521 | 6541 |
| HIV_COH__26 | Reverse | gi\|870833055\|gb\|KP178441.1\| | 479 | 452 |  | 1268 | 1241 |
| HIV_COH__27 | Reverse | gi\|870833239\|gb\|KP178445.1\| | 515 | 490 |  | 1313 | 1288 |
| HIV_COH__28 | Reverse | gi\|869135368\|gb\|KP718936.1\| | 8698 | 8681 |  | 100 | 83 |
| HIV_COH__29 | Forward | gi\|883744188\|gb\|KT008660.1\| | 1992 | 2012 |  | 8246 | 8266 |
| HIV_COH__30 | Reverse | gi\|869135526\|gb\|KR822834.1\| | 4354 | 4333 |  | 5222 | 5201 |
| HIV_COH__31 | Forward | gi\|887517014\|gb\|KT074935.1\| | 3197 | 3208 |  | 998 | 1009 |
| HIV_COH__32 | Forward | gi\|870833348\|gb\|KP178449.1\| | 1908 | 1927 |  | 4058 | 4077 |
| HIV_COH__33 | Forward | gi\|1190565781\|gb\|KY989953.1 \| | 6900 | 6921 |  | 7360 | 7381 |
| HIV_COH__34 | Forward | gi\|1190565827\|gb\|KY989956.1\| | 6950 | 6958 |  | 7522 | 7540 |
| HIV_COH__35 | Forward | gi\|869135503\|gb\|KR822832.1\| | 4916 | 4932 |  | 5520 | 5536 |
| HIV_COH__36 | Forward | gi\|869135478\|gb\|KR822830.1\| | 4796 | 4817 |  | 5438 | 5459 |
| HIV_COH__37 | Forward | gi\|871044527\|gb\|KP873164.1\| | 5077 | 5095 |  | 5563 | 5581 |
| HIV_COH__38 | Reverse | gi\|883744188\|gb\|KT008660.1\| | 246 | 227 |  | 6473 | 6454 |
| HIV_COH__39 | Reverse | gi\|658309113\|emb\|HG327288.1\| | 182 | 161 |  | 6394 | 6382 |
| HIV_COH__40 | Reverse | gi\|887517014\|gb\|KT074935.1\| | 5535 | 5514 |  | 6357 | 6336 |
| HIV_COH__41 | Forward | gi\|887517014\|gb\|KT074935.1\| | 5520 | 5538 |  | 6342 | 6360 |
| HIV_COH__42 | Forward | gi\|871044527\|gb\|KP873164.1\| | 6038 | 6058 |  | 6521 | 6541 |
| HIV_COH__43 | Reverse | gi\|870833055\|gb\|KP178441.1\| | 479 | 452 |  | 1268 | 1241 |
| HIV_COH__44 | Reverse | gi\|870833239\|gb\|KP178445.1\| | 515 | 490 |  | 1313 | 1288 |
| HIV_COH__45 | Reverse | gi\|869135368\|gb\|KP718936.1\| | 8698 | 8681 |  | 100 | 83 |
| HIV_COH__46 | Forward | gi\|883744188\|gb\|KT008660.1\| | 1992 | 2012 |  | 8246 | 8266 |
| HIV_COH__47 | Reverse | gi\|869135526\|gb\|KR822834.1\| | 4354 | 4333 |  | 5222 | 5201 |
| HIV_COH__48 | Forward | gi\|887517014\|gb\|KT074935.1\| | 3197 | 3208 |  | 998 | 1009 |
| HIV_COH__49 | Forward | gi\|870833348\|gb\|KP178449.1\| | 1908 | 1927 |  | 4058 | 4077 |
| HIV_COH__50 | Reverse | gi\|887517014\|gb\|KT074935.1\| | 4130 | 4113 |  | 4948 | 4931 |
| HIV_COH__51 | Forward | gi\|887517014\|gb\|KT074935.1\| | 3253 | 3263 |  | 4071 | 4081 |

Table S2 - *continued*

| Name | Orientation | Best hit in GenBank | | |  | Coordinates in HXB2 strain (K03455.1) | |
| --- | --- | --- | --- | --- | --- | --- | --- |
|  |  | Identification | Start coordinate | End coordinate |  | Start | End |
| HIV_COH__52 | Forward | gi\|887517014\|gb\|KT074935.1\| | 3334 | 3353 |  | 4152 | 4171 |
| HIV_COH__53 | Reverse | gi\|887517014\|gb\|KT074935.1\| | 4269 | 4250 |  | 5087 | 5068 |
| HIV_COH__54 | Forward | gi\|870833055\|gb\|KP178441.1\| | 452 | 479 |  | 1241 | 1268 |
| HIV_COH__55 | Forward | gi\|870833239\|gb\|KP178445.1\| | 490 | 515 |  | 1288 | 1313 |
| HIV_COH__56 | Forward | gi\|145207136\|gb\|EF125598.1\| | 18 | 34 |  | 2044 | 2055 |
| HIV_COH__57 | Forward | gi\|871044527\|gb\|KP873164.1\| | 1330 | 1355 |  | 1810 | 1835 |
| HIV_COH__58 | Forward | gi\|870833239\|gb\|KP178445.1\| | 485 | 507 |  | 1283 | 1305 |
| HIV_COH__59 | Forward | gi\|870833239\|gb\|KP178445.1\| | 499 | 520 |  | 1297 | 1318 |
| HIV_COH__60 | Reverse | gi\|874510212\|gb\|KP877228.1\| | 286 | 266 |  | 2538 | 2518 |
| HIV_COH__61 | Reverse | gi\|887517014\|gb\|KT074935.1\| | 1695 | 1673 |  | 2513 | 2491 |
| HIV_COH__62 | Forward | gi\|459650425\|gb\|KC312330.1\| | 1669 | 1688 |  | 2260 | 2270 |
| HIV_COH__63 | Forward | gi\|887517014\|gb\|KT074935.1\| | 1289 | 1308 |  | 2095 | 2114 |
| HIV_COH__64 | Forward | gi\|871044527\|gb\|KP873164.1\| | 922 | 942 |  | 1402 | 1422 |
| HIV_COH__65 | Forward | gi\|862573490\|emb\|HG421577.1\| | 77 | 96 |  | 2160 | 2179 |
| HIV_COH__66 | Reverse | gi\|874510014\|gb\|KP877129.1\| | 213 | 198 |  | 2465 | 2454 |
| HIV_COH__67 | Reverse | gi\|877796755\|emb\|LM654298.1\| | 66 | 46 |  | 2270 | 2250 |
| HIV_COH__68 | Forward | gi\|870833515\|gb\|KP178456.1\| | 59 | 80 |  | 7060 | 7081 |
| HIV_COH__69 | Forward | gi\|883744194\|gb\|KT008663.1\| | 1702 | 1721 |  | 7947 | 7966 |
| HIV_COH__70 | Forward | gi\|883744194\|gb\|KT008663.1\| | 1105 | 1124 |  | 7347 | 7363 |
| HIV_COH__71 | Forward | gi\|883744194\|gb\|KT008663.1\| | 1585 | 1602 |  | 7830 | 7847 |
| HIV_COH__72 | Reverse | gi\|869135240\|gb\|KP718925.1\| | 8336 | 8317 |  | 8899 | 8887 |
| HIV_COH__73 | Forward | gi\|869135380\|gb\|KP718937.1\| | 8634 | 8656 |  | 36 | 55 |
| HIV_COH__74 | Forward | gi\|869135368\|gb\|KP718936.1\| | 8673 | 8694 |  | 75 | 96 |
| HIV_COH__75 | Reverse | gi\|870831950\|gb\|KP178420.1\| | 792 | 773 |  | 781 | 762 |
| HIV_COH__76 | Reverse | gi\|870831950\|gb\|KP178420.1\| | 5643 | 5623 |  | 5662 | 5642 |
| HIV_COH__77 | Forward | gi\|870831950\|gb\|KP178420.1\| | 5160 | 5173 |  | 5164 | 5177 |
| HIV_COH__78 | Reverse | gi\|870831950\|gb\|KP178420.1\| | 5173 | 5158 |  | 5177 | 5162 |
| HIV_COH__79 | Reverse | gi\|871044527\|gb\|KP873164.1\| | 5107 | 5090 |  | 5593 | 5583 |
| HIV_COH__80 | Reverse | gi\|871044527\|gb\|KP873164.1\| | 5095 | 5077 |  | 5578 | 5563 |
| HIV_COH__81 | Forward | gi\|870833116\|gb\|KP178443.1\| | 399 | 422 |  | 1173 | 1196 |
| HIV_COH__82 | Forward | gi\|874510348\|gb\|KP877296.1\| | 682 | 701 |  | 2934 | 2953 |
| HIV_COH__83 | Reverse | gi\|874510356\|gb\|KP877300.1\| | 808 | 789 |  | 3060 | 3041 |
| HIV_COH__84 | Reverse | gi\|874507180\|gb\|KR182473.1\| | 1160 | 1139 |  | 7381 | 7360 |
| HIV_COH__85 | Forward | gi\|887517014\|gb\|KT074935.1\| | 4231 | 4250 |  | 5049 | 5068 |
| HIV_COH__86 | Forward | gi\|869135113\|gb\|KP718914.1\| | 4638 | 4657 |  | 5270 | 5289 |
| HIV_COH__87 | Forward | gi\|869135503\|gb\|KR822832.1\| | 4913 | 4932 |  | 5517 | 5536 |
| HIV_COH__88 | Forward | gi\|693792547\|gb\|KM217867.1\| | 37 | 56 |  | 5671 | 5686 |
| HIV_COH__89 | Reverse | gi\|612529827\|gb\|KJ017738.1\| | 97 | 78 |  | 5225 | 5214 |
| HIV_COH__90 | Reverse | gi\|761545410\|gb\|KP411830.1\| | 4674 | 4656 |  | 5463 | 5445 |
| HIV_COH__91 | Reverse | gi\|869135206\|gb\|KP718922.1\| | 5110 | 5091 |  | 5723 | 5704 |
| HIV_COH__92 | Reverse | gi\|761545386\|gb\|KP411827.1\| | 5096 | 5077 |  | 5868 | 5849 |
| HIV_COH__93 | Forward | gi\|877797056\|emb\|LM654303.1\| | 867 | 887 |  | 5069 | 5089 |
| HIV_COH__94 | Forward | gi\|256674250\|gb\|FJ641184.1\| | 380 | 401 |  | 5323 | 5342 |
| HIV_COH__95 | Forward | gi\|822094503\|gb\|KP455640.1\| | 4699 | 4720 |  | 5546 | 5567 |
| HIV_COH__96 | Forward | gi\|693800206\|gb\|KM218259.1\| | 790 | 809 |  | 5710 | 5729 |
| HIV_COH__97 | Reverse | gi\|869135526\|gb\|KR822834.1\| | 4353 | 4334 |  | 5221 | 5202 |
| HIV_COH__98 | Reverse | gi\|437191028\|gb\|JX149433.1\| | 415 | 396 |  | 5453 | 5437 |
| HIV_COH__99 | Reverse | gi\|672919174\|gb\|KJ849816.1\| | 5167 | 5148 |  | 5686 | 5676 |
| HIV_COH__100 | Reverse | gi\|693799192\|gb\|KM218209.1\| | 109 | 90 |  | 5839 | 5820 |
| HIV_COH__101 | Forward | gi\|7416477\|dbj\|AB034427.1\| | 175 | 197 |  | 6013 | 6027 |
| HIV_COH__102 | Forward | gi\|887517014\|gb\|KT074935.1\| | 5390 | 5409 |  | 6209 | 6228 |
| HIV_COH__103 | Forward | gi\|855687902\|gb\|KT152843.1\| | 5794 | 5813 |  | 6459 | 6473 |
| HIV_COH__104 | Reverse | gi\|693799192\|gb\|KM218209.1\| | 500 | 480 |  | 6228 | 6210 |
| HIV_COH__105 | Reverse | gi\|846121816\|gb\|KR861266.1\| | 483 | 464 |  | 6446 | 6431 |
| HIV_COH__106 | Reverse | gi\|769347149\|gb\|KJ952265.1\| | 2463 | 2446 |  | 1167 | 1160 |

Table S2 - *continued*

| Name | Orientation | Best hit in GenBank | | |  | Coordinates in HXB2 strain (K03455.1) | |
| --- | --- | --- | --- | --- | --- | --- | --- |
|  |  | Identification | Start coordinate | End coordinate |  | Start | End |
| HIV_COH__107 | Forward | gi\|761545386\|gb\|KP411827.1\| | 5250 | 5276 |  | 6022 | 6048 |
| HIV_COH__108 | Forward | gi\|846575165\|gb\|KP693383.1\| | 33 | 49 |  | 5039 | 5047 |
| HIV_COH__109 | Forward | gi\|354720562\|gb\|JF680917.1\| | 248 | 267 |  | 6486 | 6497 |
| HIV_COH__110 | Reverse | gi\|769793172\|gb\|KM081847.1\| | 155 | 129 |  | 6216 | 6190 |
| HIV_COH__111 | Reverse | gi\|874507180\|gb\|KR182473.1\| | 152 | 132 |  | 6379 | 6363 |
| HIV_COH__112 | Reverse | gi\|883744194\|gb\|KT008663.1\| | 376 | 354 |  | 6603 | 6581 |
| HIV_COH__113 | Forward | gi\|887517014\|gb\|KT074935.1\| | 6160 | 6178 |  | 6955 | 6973 |
| HIV_COH__114 | Forward | gi\|874507180\|gb\|KR182473.1\| | 793 | 811 |  | 7008 | 7026 |
| HIV_COH__115 | Reverse | gi\|887517014\|gb\|KT074935.1\| | 6724 | 6706 |  | 7540 | 7522 |
| HIV_COH__116 | Reverse | gi\|883744194\|gb\|KT008663.1\| | 2341 | 2323 |  | 8586 | 8568 |
| HIV_COH__117 | Forward | gi\|874507180\|gb\|KR182473.1\| | 1562 | 1580 |  | 7795 | 7813 |
| HIV_COH__118 | Reverse | gi\|887517014\|gb\|KT074935.1\| | 705 | 686 |  | 1505 | 1486 |
| HIV_COH__119 | Reverse | gi\|874510206\|gb\|KP877225.1\| | 1298 | 1279 |  | 3550 | 3531 |
| HIV_COH__120 | Forward | gi\|824907897\|gb\|KR781669.1\| | 1186 | 1203 |  | 1976 | 1993 |
| HIV_COH__121 | Forward | gi\|870833239\|gb\|KP178445.1\| | 1213 | 1232 |  | 2011 | 2030 |
| IV_COH__122 | Forward | gi\|870833239\|gb\|KP178445.1\| | 1221 | 1240 |  | 2019 | 2038 |
| HIV_COH__123 | Forward | gi\|870833116\|gb\|KP178443.1\| | 1443 | 1462 |  | 2217 | 2236 |
| HIV_COH__124 | Forward | gi\|558756595\|dbj\|AB873982.1\| | 166 | 185 |  | 2250 | 2269 |
| HIV_COH__125 | Forward | gi\|874510180\|gb\|KP877212.1\| | 48 | 67 |  | 2300 | 2319 |
| HIV_COH__126 | Forward | gi\|874510156\|gb\|KP877200.1\| | 103 | 123 |  | 2355 | 2375 |
| HIV_COH__127 | Forward | gi\|887517014\|gb\|KT074935.1\| | 1575 | 1594 |  | 2393 | 2412 |
| HIV_COH__128 | Forward | gi\|814946187\|gb\|KP250807.1\| | 165 | 190 |  | 2432 | 2457 |
| HIV_COH__129 | Forward | gi\|887517014\|gb\|KT074935.1\| | 1673 | 1695 |  | 2491 | 2513 |
| HIV_COH__130 | Forward | gi\|874510212\|gb\|KP877228.1\| | 269 | 290 |  | 2521 | 2542 |
| HIV_COH__131 | Forward | gi\|814946209\|gb\|KP250813.1\| | 298 | 319 |  | 2550 | 2571 |
| HIV_COH__132 | Forward | gi\|874510212\|gb\|KP877228.1\| | 344 | 363 |  | 2596 | 2615 |
| HIV_COH__133 | Forward | gi\|874510156\|gb\|KP877200.1\| | 416 | 436 |  | 2668 | 2688 |
| HIV_COH__134 | Forward | gi\|887517014\|gb\|KT074935.1\| | 1871 | 1890 |  | 2689 | 2708 |
| HIV_COH__135 | Forward | gi\|874510294\|gb\|KP877269.1\| | 491 | 516 |  | 2743 | 2768 |
| HIV_COH__136 | Forward | gi\|874510358\|gb\|KP877301.1\| | 555 | 576 |  | 2807 | 2828 |
| HIV_COH__137 | Forward | gi\|874510300\|gb\|KP877272.1\| | 582 | 600 |  | 2834 | 2852 |
| HIV_COH__138 | Forward | gi\|874510210\|gb\|KP877227.1\| | 624 | 644 |  | 2876 | 2896 |
| HIV_COH__139 | Forward | gi\|874510206\|gb\|KP877225.1\| | 699 | 724 |  | 2951 | 2976 |
| HIV_COH__140 | Forward | gi\|887517014\|gb\|KT074935.1\| | 2180 | 2199 |  | 2998 | 3017 |
| HIV_COH__141 | Forward | gi\|874510258\|gb\|KP877251.1\| | 779 | 805 |  | 3031 | 3057 |
| HIV_COH__142 | Forward | gi\|874510342\|gb\|KP877293.1\| | 848 | 872 |  | 3100 | 3124 |
| HIV_COH__143 | Forward | gi\|874510212\|gb\|KP877228.1\| | 882 | 901 |  | 3134 | 3153 |
| HIV_COH__144 | Forward | gi\|874510082\|gb\|KP877163.1\| | 919 | 938 |  | 3171 | 3190 |
| HIV_COH__145 | Forward | gi\|874510212\|gb\|KP877228.1\| | 973 | 992 |  | 3225 | 3244 |
| HIV_COH__146 | Forward | gi\|874510206\|gb\|KP877225.1\| | 1023 | 1043 |  | 3275 | 3295 |
| HIV_COH__147 | Forward | gi\|874510196\|gb\|KP877220.1\| | 1069 | 1091 |  | 3321 | 3343 |
| HIV_COH__148 | Forward | gi\|874510172\|gb\|KP877208.1\| | 1101 | 1123 |  | 3353 | 3375 |
| HIV_COH__149 | Forward | gi\|844288908\|gb\|KR861122.1\| | 1900 | 1921 |  | 3393 | 3414 |
| HIV_COH__150 | Forward | gi\|874510212\|gb\|KP877228.1\| | 1193 | 1214 |  | 3445 | 3466 |
| HIV_COH__151 | Forward | gi\|874510210\|gb\|KP877227.1\| | 1242 | 1264 |  | 3494 | 3516 |
| HIV_COH__152 | Forward | gi\|870833116\|gb\|KP178443.1\| | 1269 | 1290 |  | 2043 | 2064 |
| HIV_COH__153 | Forward | gi\|693446106\|gb\|KJ869543.1\| | 1302 | 1322 |  | 2094 | 2114 |
| HIV_COH__154 | Forward | gi\|870833239\|gb\|KP178445.1\| | 1319 | 1338 |  | 2117 | 2136 |
| HIV_COH__155 | Forward | gi\|862573490\|emb\|HG421577.1\| | 71 | 89 |  | 2154 | 2172 |
| HIV_COH__156 | Forward | gi\|672611500\|gb\|KJ925006.1\| | 2210 | 2229 |  | 2174 | 2193 |
| HIV_COH__157 | Forward | gi\|870832714\|gb\|KP178423.1\| | 1407 | 1426 |  | 2193 | 2212 |
| HIV_COH__158 | Reverse | gi\|870833146\|gb\|KP178444.1\| | 1260 | 1237 |  | 2082 | 2059 |
| HIV_COH__159 | Reverse | gi\|813549609\|gb\|KP995629.1\| | 21 | 2 |  | 2273 | 2254 |
| HIV_COH__160 | Reverse | gi\|874510372\|gb\|KP877308.1\| | 95 | 72 |  | 2347 | 2324 |
| HIV_COH__161 | Reverse | gi\|874510206\|gb\|KP877225.1\| | 120 | 100 |  | 2372 | 2352 |

Table S2 - *continued*

| Name | Orientation | Best hit in GenBank | | |  | Coordinates in HXB2 strain (K03455.1) | |
| --- | --- | --- | --- | --- | --- | --- | --- |
|  |  | Identification | Start coordinate | End coordinate |  | Start | End |
| HIV_COH__162 | Forward | gi\|887517014\|gb\|KT074935.1\| | 1222 | 1244 |  | 2028 | 2050 |
| HIV_COH__163 | Forward | gi\|887517014\|gb\|KT074935.1\| | 1222 | 1244 |  | 2028 | 2050 |
| HIV_COH__164 | Reverse | gi\|814946187\|gb\|KP250807.1\| | 194 | 168 |  | 2461 | 2435 |
| HIV_COH__165 | Reverse | gi\|887517014\|gb\|KT074935.1\| | 1695 | 1673 |  | 2513 | 2491 |
| HIV_COH__166 | Reverse | gi\|874510212\|gb\|KP877228.1\| | 286 | 266 |  | 2538 | 2518 |
| HIV_COH__167 | Reverse | gi\|814946232\|gb\|KP250819.1\| | 314 | 294 |  | 2566 | 2546 |
| HIV_COH__168 | Reverse | gi\|887517014\|gb\|KT074935.1\| | 1783 | 1762 |  | 2601 | 2580 |
| HIV_COH__169 | Reverse | gi\|870832919\|gb\|KP178437.1\| | 1316 | 1297 |  | 2105 | 2086 |
| HIV_COH__170 | Reverse | gi\|874510212\|gb\|KP877228.1\| | 372 | 353 |  | 2624 | 2605 |
| HIV_COH__171 | Reverse | gi\|887517014\|gb\|KT074935.1\| | 1889 | 1872 |  | 2707 | 2690 |
| HIV_COH__172 | Reverse | gi\|874510370\|gb\|KP877307.1\| | 475 | 452 |  | 2727 | 2704 |
| HIV_COH__173 | Reverse | gi\|874510300\|gb\|KP877272.1\| | 600 | 582 |  | 2852 | 2834 |
| HIV_COH__174 | Reverse | gi\|874510212\|gb\|KP877228.1\| | 639 | 620 |  | 2891 | 2872 |
| HIV_COH__175 | Reverse | gi\|874510210\|gb\|KP877227.1\| | 654 | 632 |  | 2906 | 2884 |
| HIV_COH__176 | Reverse | gi\|874510370\|gb\|KP877307.1\| | 714 | 688 |  | 2966 | 2940 |
| HIV_COH__177 | Reverse | gi\|874510212\|gb\|KP877228.1\| | 752 | 733 |  | 3004 | 2985 |
| HIV_COH__178 | Reverse | gi\|869135322\|gb\|KP718932.1\| | 1454 | 1435 |  | 2117 | 2098 |
| HIV_COH__179 | Reverse | gi\|874510338\|gb\|KP877291.1\| | 780 | 760 |  | 3032 | 3012 |
| HIV_COH__180 | Reverse | gi\|874510236\|gb\|KP877240.1\| | 824 | 801 |  | 3076 | 3053 |
| HIV_COH__181 | Reverse | gi\|874510342\|gb\|KP877293.1\| | 869 | 845 |  | 3121 | 3097 |
| HIV_COH__182 | Reverse | gi\|874510352\|gb\|KP877298.1\| | 890 | 870 |  | 3142 | 3122 |
| HIV_COH__183 | Reverse | gi\|874510082\|gb\|KP877163.1\| | 938 | 919 |  | 3190 | 3171 |
| HIV_COH__184 | Reverse | gi\|874510368\|gb\|KP877306.1\| | 987 | 968 |  | 3239 | 3220 |
| HIV_COH__185 | Reverse | gi\|874510366\|gb\|KP877305.1\| | 1018 | 996 |  | 3270 | 3248 |
| HIV_COH__186 | Reverse | gi\|874510206\|gb\|KP877225.1\| | 1048 | 1029 |  | 3300 | 3281 |
| HIV_COH__187 | Reverse | gi\|874510372\|gb\|KP877308.1\| | 1064 | 1043 |  | 3316 | 3295 |
| HIV_COH__188 | Reverse | gi\|874510204\|gb\|KP877224.1\| | 1117 | 1097 |  | 3369 | 3349 |
| HIV_COH__189 | Reverse | gi\|870833146\|gb\|KP178444.1\| | 1331 | 1312 |  | 2153 | 2134 |
| HIV_COH__190 | Reverse | gi\|874510032\|gb\|KP877138.1\| | 1161 | 1140 |  | 3413 | 3392 |
| HIV_COH__191 | Reverse | gi\|874510212\|gb\|KP877228.1\| | 1208 | 1187 |  | 3460 | 3439 |
| HIV_COH__192 | Reverse | gi\|874510134\|gb\|KP877189.1\| | 1225 | 1203 |  | 3477 | 3455 |
| HIV_COH__193 | Reverse | gi\|874510206\|gb\|KP877225.1\| | 1300 | 1281 |  | 3552 | 3533 |
| HIV_COH__194 | Forward | gi\|887517014\|gb\|KT074935.1\| | 1222 | 1244 |  | 2028 | 2050 |
| HIV_COH__195 | Reverse | gi\|767577186\|gb\|KP113466.1\| | 53 | 34 |  | 2188 | 2169 |
| HIV_COH__196 | Reverse | gi\|870832714\|gb\|KP178423.1\| | 1426 | 1407 |  | 2212 | 2193 |
| HIV_COH__197 | Reverse | gi\|870833116\|gb\|KP178443.1\| | 1453 | 1434 |  | 2227 | 2208 |
| HIV_COH__198 | Reverse | gi\|542102483\|gb\|KF526193.1\| | 1609 | 1588 |  | 2269 | 2248 |
| HIV_COH__199 | Forward | gi\|874510372\|gb\|KP877308.1\| | 228 | 247 |  | 2480 | 2499 |
| HIV_COH__200 | Reverse | gi\|869135515\|gb\|KR822833.1\| | 2763 | 2742 |  | 3420 | 3399 |
| HIV_COH__201 | Forward | gi\|887517014\|gb\|KT074935.1\| | 1222 | 1244 |  | 2028 | 2050 |
| HIV_COH__202 | Forward | gi\|869135537\|gb\|KR822835.1\| | 7921 | 7937 |  | 8492 | 8508 |
| HIV_COH__203 | Forward | gi\|883744188\|gb\|KT008660.1\| | 2252 | 2270 |  | 8506 | 8524 |
| HIV_COH__204 | Forward | gi\|887517014\|gb\|KT074935.1\| | 7718 | 7739 |  | 8522 | 8543 |
| HIV_COH__205 | Forward | gi\|869135515\|gb\|KR822833.1\| | 8147 | 8167 |  | 8789 | 8809 |
| HIV_COH__206 | Reverse | gi\|869135515\|gb\|KR822833.1\| | 8872 | 8851 |  | 436 | 423 |
| HIV_COH__207 | Reverse | gi\|883744188\|gb\|KT008660.1\| | 2270 | 2252 |  | 8524 | 8506 |
| HIV_COH__208 | Reverse | gi\|874506439\|gb\|KR182324.1\| | 2328 | 2307 |  | 8543 | 8522 |
| HIV_COH__209 | Reverse | gi\|44888946\|gb\|AY452651.1\| | 431 | 411 |  | 435 | 423 |
| HIV_COH__210 | Reverse | gi\|874510288\|gb\|KP877266.1\| | 1067 | 1048 |  | 3319 | 3300 |
| HIV_COH__211 | Reverse | gi\|874510212\|gb\|KP877228.1\| | 1210 | 1189 |  | 3462 | 3441 |
| HIV_COH__212 | Reverse | gi\|874510366\|gb\|KP877305.1\| | 1287 | 1257 |  | 3539 | 3509 |
| HIV_COH__213 | Forward | gi\|869135515\|gb\|KR822833.1\| | 8954 | 8975 |  | 518 | 539 |
| HIV_COH__214 | Reverse | gi\|877797352\|emb\|LM654310.1\| | 630 | 611 |  | 4826 | 4807 |
| HIV_COH__215 | Forward | gi\|877797352\|emb\|LM654310.1\| | 611 | 630 |  | 4807 | 4826 |
| HIV_COH__216 | Reverse | gi\|761545404\|gb\|KP411829.1\| | 4854 | 4831 |  | 5649 | 5626 |
| HIV_COH__217 | Reverse | gi\|871044527\|gb\|KP873164.1\| | 5107 | 5089 |  | 5593 | 5575 |

Table S2 - *continued*

| Name | Orientation | Best hit in GenBank | | |  | Coordinates in HXB2 strain (K03455.1) | |
| --- | --- | --- | --- | --- | --- | --- | --- |
|  |  | Identification | Start coordinate | End coordinate |  | Start | End |
| HIV_COH__218 | Forward | gi\|869135537\|gb\|KR822835.1\| | 5396 | 5416 |  | 5962 | 5982 |
| HIV_COH__219 | Reverse | gi\|874506736\|gb\|KR182384.1\| | 611 | 589 |  | 6835 | 6813 |
| HIV_COH__220 | Reverse | gi\|761631427\|gb\|KP411823.1\| | 5225 | 5211 |  | 6027 | 6013 |
| HIV_COH__221 | Reverse | gi\|846122140\|gb\|KR861332.1\| | 39 | 20 |  | 6005 | 5986 |
| HIV_COH__222 | Forward | gi\|887517014\|gb\|KT074935.1\| | 5003 | 5023 |  | 5822 | 5842 |
| HIV_COH__223 | Reverse | gi\|887517014\|gb\|KT074935.1\| | 5534 | 5512 |  | 6356 | 6334 |

**Table S3.** PCR primers used to amplify and sequence the human gene encoding APOBEC3G

| Primer identification | Sequence (5’-3’) |
| --- | --- |
| A3G_DNA_FR1_Fw1 | AGGGAGGGCTGTCCTAAAAC |
| A3G_DNA_FR1_R1 | CATCAAAGCCCTGTGAACAA |
| A3G_DNA_FR1_Fw2 | AAAACCAGAAGCTTGGAGCA |
| A3G_DNA_FR1_R2 | GAAGCACACAGAGGGAGCAC |
| A3G_DNA_FR2_Fw1 | GACGGAATTTCGCTCTTGTC |
| A3G_DNA_FR2_R | GCAGCTACTTGCGAGTCTGA1 |
| A3G_DNA_FR2_Fw2 | CCAGGGGGCTGTGTTTAGT |
| A3G_DNA_FR2_R2 | CCTTTTGTGCAGAGGCAAAC |
| A3G_DNA_FR34_Fw1 | AAGGAGCTTCAATGGCAAGA |
| A3G_DNA_FR34_R1 | TCGGTTATGCCAGGGAGAT |
| A3G_DNA_FR34_Fw2 | TGGCAAGATCTCCTGGACTC |
| A3G_DNA_FR34_R2 | AGAAGGGGCCTCTCACTCAC |
| A3G_DNA_FR34_Fw3 | TGCTCCTCTCCCAGGTGTAT |
| A3G_DNA_FR34_R3 | GAGGATGAGCAGGAGGTGAG |
| A3G_DNA_FR5_Fw1 | GGGAAGAAGACCCAGCTGTA |
| A3G_DNA_FR5_R1 | AGTGGTGTTAGCCCCTGAGA |
| A3G_DNA_FR5_Fw2 | GGAGACCCTGACAAGGCTTAG |
| A3G_DNA_FR5_R2 | CCACAGGTACCACCCAGAAC |
| A3G_DNA_FR5_Fw3 | GGTTTGGAGGCTCTAGCAAGT |
| A3G_DNA_FR6_Fw1 | GAGACCAGCCTGAACCACAT |
| A3G_DNA_FR6_R1 | CCCAAGTTCTGCTTCCACTT |
| A3G_DNA_FR6_Fw2 | CCTGGGCAACAAGAGTGAA |
| A3G_DNA_FR6_R2 | CTCCCCAGGTCCACATCTC |
| A3G_DNA_FR6_Fw3 | CCTCATGGCTTGCTTTCTTT |
| A3G_DNA_FR7_R1 | AGAAAGGGAGGCTGTGAGAA |
| A3G_DNA_FR7_R2 | GAAAGGGAGGGACAGAGACA |
| A3G_DNA_FR7_R3 | CACTGGGCAGAGGGAAGA |
| A3G_DNA_FR1_R1-Frw | TTGTTCACAGGGCTTTGATG |
| A3G_DNA_FR1_R2-Frw | GTGCTCCCTCTGTGTGCTTC |
| A3G_DNA_FR2_Fw1-Rev | GACAAGAGCGAAATTCCGTC |
| A3G_DNA_FR2_Fw2-Rev | ACTAAACACAGCCCCCTGG |
| A3G_DNA_FR2_R1-Frw | TCAGACTCGCAAGTAGCTGC |
| A3G_DNA_FR2_R2-Frw | GTTTGCCTCTGCACAAAAGG |
| A3G_DNA_FR34_Fw1-Rev | TCTTGCCATTGAAGCTCCTT |
| A3G_DNA_FR34_Fw2-Rev | GAGTCCAGGAGATCTTGCCA |
| A3G_DNA_FR34_R1-Frw | ATCTCCCTGGCATAACCGA |
| A3G_DNA_FR34_R2-Fwr | GTGAGTGAGAGGCCCCTTCT |
| A3G_DNA_FR5_Fw1-Rev | TACAGCTGGGTCTTCTTCCC |
| A3G_DNA_FR5_Fw2-Rev | CTAAGCCTTGTCAGGGTCTCC |
| A3G_DNA_FR5_R1-Frw | TCTCAGGGGCTAACACCACT |
| A3G_DNA_FR5_R2-Frw | GTTCTGGGTGGTACCTGTGG |
| A3G_DNA_FR6_Fw1-Rev | ATGTGGTTCAGGCTGGTCTC |
| A3G_DNA_FR6_Fw2-Rev | TTCACTCTTGTTGCCCAGG |
| A3G_DNA_FR78_Fw1 | GCTGGAAGTGGAAGCAGAAC |
| A3G_DNA_FR78_R1 | GATTCAAATTGTCGCTGGATTA |
| A3G_DNA_FR78_Fw2 | AGGAAAGAAGTGCCTGATGAA |
| A3G_DNA_FR78_R2 | TGTCGCTGGATTAGTAGGTTTG |
| A3G_DNA_FR78_R3 | CATTGCTTTGCTGGTGTCTG |
| A3G_DNA_INT56A_Fw1 | CAGAGGACGGCATGAGACTT |
| A3G_DNA_INT56A_R1 | TCCCATCCCACTGGTTAGAC |
| A3G_DNA_INT56A_Fw2 | GAGCGCATGCACAATGAC |
| A3G_DNA_INT56A_R2 | CAGGGCCCATATTAAGCTCAT |
| A3G_DNA_INT56B_Fw1 | GGCCAGCATTTGAAGAAGAG |
| A3G_DNA_INT56B_R1 | GGCCGAGGGAATTTTTAGTG |
| A3G_DNA_INT56B_Fw2 | CCAAATTCTGCTTGGCTCTG |
| A3G_DNA_INT56B_R2 | GAGGGACAGCCTGGATCA |
